# Mutation in the TRKB cholesterol recognition site that blocks antidepressant binding does not influence the basal or BDNF-stimulated activation of TRKB

**DOI:** 10.1101/2022.08.26.505413

**Authors:** Caroline Biojone, Cecilia Cannarozzo, Nina Seiffert, Cassiano RAF Diniz, Cecilia A Brunello, Eero Castrén, Plinio Casarotto

## Abstract

Brain-derived neurotrophic factor (BDNF) acting upon its receptor Neurotrophic tyrosine kinase receptor 2 (NTRK2, TRKB) plays a central role in the development and maintenance of synaptic function and activity- or drug-induced plasticity. TRKB possesses an inverted cholesterol-recognition and alignment consensus sequence (CARC), suggesting this receptor can act as a cholesterol sensor. We have recently shown that antidepressants drugs directly bind to the CARC domain of TRKB dimers, and that this binding as well as biochemical and behavioral responses to antidepressants are lost with a mutation in the TRKB CARC motif (Y433F). However, it is not clear if this mutation can also compromise the receptor function and lead to behavioral alterations. Here, we observed that Y433F mutation does not alter BDNF binding to TRKB, or BDNF-induced dimerization of TRKB. In this line, primary cultures from embryos of heterozygous Y433F mutant mice (hTRKB.Y433F) are responsive to BDNF-induced activation of TRKB, and samples from adult mice do not show any difference on TRKB activation compared to wild-type littermates (TRKB.wt). The behavioral phenotype of hTRKB.Y433F mice is indistinguishable from the wild-type mice in cued fear conditioning, contextual discrimination task or the elevated plus maze, whereas mice heterozygous to BDNF null allele show a phenotype in context discrimination task. Taken together, our results indicate that Y433F mutation in the TRKB CARC motif does not show signs of loss-of-function of BDNF responses, while antidepressant binding to TRKB and responses to antidepressants are lost in Y433F mutants, making them an interesting mouse model for antidepressant research.

## Introduction

Brain-derived neurotrophic factor (BDNF) is a critical regulator of neuronal connectivity and plasticity, and these effects are mediated by binding of BDNF to neurotrophic tyrosine kinase receptor 2 (NTRK2, TRKB) (Bothwell, 2016). Neuronal plasticity is compromised in several brain disorders and promotion of plasticity would be highly useful in the prevention and treatment of, as well as in recovery and rehabilitation after several neurological and psychiatric disorders (Castrén and Antila, 2017). Indeed, there is a widespread consensus that BDNF-TRKB mediated plasticity is a critical mediator of the effects of antidepressant drugs and recent evidence indicates that TRKB is the direct site of action of antidepressant binding (Casarotto et al., 2021).

Cholesterol is another important regulator of synaptic maturation and plasticity (Mauch et al., 2001; Martín et al., 2014). Dietary cholesterol does not pass into the brain and the nervous system is dependent on locally produced cholesterol (Martín et al., 2014). The majority of neuronal cholesterol is produced by astrocytes that provide neurons with cholesterol through an ApoE-mediated transport (Pfrieger, 2003; Martín et al., 2014). Cholesterol is an integral component of plasma membrane and cholesterol concentration within different plasma membrane compartments varies from low levels to up to 50 mol% in synaptic membranes (Ikonen, 2008; Tulodziecka et al., 2016). Cholesterol concentrations in synaptic membranes increase during postnatal development, along with synaptic maturation (Tulodziecka et al., 2016) and in the absence of cholesterol, synapses fail to mature and synaptic signaling is compromised (Mauch et al., 2001).

Several lines of evidence suggest that TRKB is a major target through which the effects of cholesterol on neuronal plasticity are mediated. BDNF signaling increases the cholesterol production in neurons (Suzuki et al., 2013; Zonta and Minichiello, 2013) and TRKB signaling is regulated by cholesterol (Suzuki et al., 2004; Pereira and Chao, 2007). We have recently shown that TRKB, but not other TRK-family members, possesses an inverted cholesterol recognition domain (CARC) within the transmembrane region (Cannarozzo et al., 2021; Casarotto et al., 2021).

Cholesterol apparently influences TRKB function through at least two mechanisms. First, through direct interaction with TRKB through the CARC domain, as a tyrosine to phenylalanine point mutation in the position 433 (TRKB.Y433F), which is considered a critical component of the CARC domain (Fantini et al., 2019), blocks the facilitatory effects of cholesterol on TRKB signaling (Casarotto et al., 2021). Second, and our data suggest that this is the main effect, cholesterol regulates the configuration of TRKB transmembrane domains (TMD) though its effects on membrane thickness: membranes rich in cholesterol, such as synaptic membranes, are thicker, which influences the positioning of the TRKB TMDs. TRKB TMD possesses a ^439^AXXXG^443^ dimerization motif, through which the TMDs of TRKB dimers crisscross each other within the plasma membrane, analogous to the EGF receptor (Arkhipov et al., 2013; Endres et al., 2013; Sinclair et al., 2018). In moderate cholesterol concentrations, TRKB assumes a crossed configuration that is competent for efficient BDNF signaling. In thick membranes, however, the crossed configuration is disrupted and TRKB is not stable, which is consistent with previous observations that TRKB is excluded from thick lipid raft membranes (Suzuki et al., 2004; Pereira and Chao, 2007). Indeed, our previous data suggest that cholesterol has a bell-shaped effect on TRKB signaling: moderate increase in plasma membrane cholesterol promotes TRB signaling, while very low and high concentrations inhibit it (Casarotto et al., 2021). This effect appears to be mostly mediated by a decreased TRKB localization at the cell surface in high cholesterol concentrations (Cannarozzo et al., 2021), which is consistent with exclusion of TRKB from synaptic membranes (Suzuki et al., 2004; Pereira and Chao, 2007).

We recently discovered that antidepressant drugs, including the typical serotonin selective reuptake inhibitors (SSRI) and tricyclic antidepressants (TCA) as well as the rapid-acting antidepressant ketamine, directly bind to the TRKB CARC domain, thereby stabilizing the signaling-competent TRKB structure in synaptic membranes and allosterically increasing BDNF signaling through TRKB (Casarotto et al., 2021). Indeed, transgenic mice heterozygous for a Y433F point mutation in the CARC domain fail to show any plasticity-promoting and antidepressant-like behavioral responses of antidepressant administration (Casarotto et al., 2021). It is unclear, however, whether the TRKB.Y433F mutation has any effects of the on basic expression or localisation of TRKB, or on the ability of BDNF to activate and signal through TRKB, and whether such effects might contribute to the inability of antidepressants to influence the mutant TRKB.

We have here investigated the effects of the TRKB.Y433F mutation on baseline activity of TRKB and on the effects of BDNF through the mutated receptor *in vitro* and *in vivo*. As our previous studies have shown that the effects of antidepressant drugs are lost in heterozygous mice with one Y433F mutant allele, we have focused on the analysis of the heterozygous mutants. We found that heterozygous TRKB.Y433F mutation (hTRKB.Y433F) has no effects on the baseline signaling of TRKB, which is consistent with our earlier observations that Y433F mutation does not influence baseline TRKB localisation or signaling (Casarotto et al., 2021). Furthermore, we found no effects of TRKB.Y433F signaling on BDNF binding or plasticity-related behavioral responses that are sensitive to BDNF. These data suggest that the hTRKB.Y433F mutation has no effect on the normal function of TRKB, but it is not sensitive to the effects of typical antidepressant drugs or ketamine.

## Material and Methods

### Animals

Male and female mice (C57BL/6JRccHsd background, Envigo-Harlan Labs, Netherlands) heterozygous to TRKB.Y433F or TRKB.wt littermates were used, the animals were bred and genotyped as described previously (Casarotto et al., 2021). For comparison, male adult mice of C57BL/6J-000664 background (from Jackson Laboratories), carrying a deletion in one of the copies of *Bdnf* gene (BDNF.het) or BDNF.wt littermates were used (Karpova et al., 2011).

The mice (maintained in the Laboratory Animal Center of the University of Helsinki) were 16-18 weeks old at the beginning of the experiments, group-housed (3-6 per cage - type ll: 552 cm2 floor area, Tecniplast, Italy) under a 12h light/dark cycle, with free access to water and food. All protocols were approved by the ethics committee for animal experimentation of Southern Finland (ESAVI/38503/2019).

### Cell culture and sample collection

Cultures of cortical cells from mouse embryos were prepared as previously described in detail (Sahu et al., 2019). Briefly, suspended cortical cells from each embryo were seeded in poly-L-lysine-coated 24-well plates (View Plate 96, PerkinElmer) at 250000 cells/well. The cells were maintained in Neurobasal medium, supplemented with B27 and left undisturbed, except for medium change (1/3 twice per week). During the culture preparation samples from each embryo were collected for genotyping as described previously (Casarotto et al., 2021).

Mouse neuroblastoma cell line N2A were cultured as described previously (Casarotto et al., 2021). Briefly, the cells were cultivated in DMEM (Lonza, cat# BE12-614Q), supplemented with 10% v/v inactivated fetal bovine serum (Invitrogen), 1% v/v L-glutamine and penicillin/streptomycin (Lonza, cat# DE17-603E), at 37°C in 5% CO_2_.

### Direct Binding Assay - DBA

The cell-free assays (binding assays) were performed in white 96-well plates as described before (Casarotto et al., 2021). The plates were precoated with anti-GFP antibodies (Abcam, #ab290, 1:1000) in carbonate buffer (pH 9.8), ON at 4°C. Following blocking with 3% BSA in PBS buffer (2 h at RT), 120 μg of total protein from each sample (of lysates from HEK293T cells transfected to overexpress GFP-TRKB.wt, GFP-TRKB.Y433F or GFP-TRKB/TRKA.TM) were added and incubated overnight at 4°C under agitation. The plates were then washed 3x with PBS buffer, and various concentrations of biotinylated BDNF (Alomone Labs, cat#B-250, 0.1 – 100 pM) was added for 1h at RT. The luminescence was determined via HRP-conjugated streptavidin (Thermo-Fisher, cat#21126, 1:10000, 1h, RT) activity reaction with ECL by a plate reader. The luminescence signal from blank wells (containing all the reagents but the sample lystates, substituted by the blocking buffer) was used as background. The specific signal was then calculated by subtracting the values of blank wells from the values of the samples with matched concentration of the biotinylated ligand.

### Enzyme-linked Immunosorbent Assay - ELISA

The levels of phosphorylated TRKB in primary cortical cultures of mouse embryos were determined by ELISA as previously described (Fred et al., 2019; Casarotto et al., 2021). Briefly, the cells were homogenized in cold NP lysis buffer (20 mM Tris-HCl, 150 mM NaCl, 50 mM NaF, 1% Nonidet-40, 10% glycerol) supplemented with 2 mM Na3VO4 and cOmplete inhibitor mix (Roche), centrifuged (4°C, 15,000 g, 10 min) and stored at -80°C. For the ELISA assay, goat anti-TRKB antibody (R&D System, #AF1494) was diluted 1:500 in carbonate buffer pH 9.7 (25 mM sodium bicarbonate, 25 Mm sodium carbonate) and coated in flat-bottom white plates (OptiPlate 96F-HB, Perkin Elmer) overnight at 4°C under agitation. Unbound antibody was then removed from the plate, and 5% BSA in PBST was incubated for 2h at RT to block nonspecific binding sites. The samples were thawed on ice, transferred to the ELISA plate, and incubated overnight under agitation at 4°C. The plates were washed 3x with PBST and incubated overnight at 4°C under agitation with rabbit anti-phosphorylated TRKB (Cell Signaling, either #4168 against pY816 or #4619 against pY515) diluted 1:1000 in 5% BSA/PBST. After a washing step (3x PBST), the plates were incubated with 1:5000 anti-rabbit HRP-conjugated tertiary antibody for 2h at RT. Plates were washed again in PBST, ECL (Thermo Fisher Scientific) was added, and the luminescence was detected by Varioskan Flash plate reader (Thermo Scientific). Unspecific signal (from wells where sample was omitted) was assessed in each plate and subtracted from samples signal. Final data was expressed as percentage of the control (vehicle treated group).

### Western blotting - WB

The levels of pTRKB and total TRKB in the prefrontal cortex of experimentally naive adult male mice was determined by WB as previously described (Rantamäki et al., 2007). Samples were homogenized as mentioned above and denatured in a 2X Laemmli buffer for 5 min at 95° C. SDS-PAGE was carried out by resolving the samples by electrophoresis in NuPAGE 4-12% Bis-Tris Protein polyacrylamide gels (#NP0323BOX, Invitrogen). After the electrophoresis, the samples were transferred to a PVDF membrane and incubated in primary antibody diluted 1:1000 in 3% BSA/TBST overnight at 4° C. The membrane was subsequently washed and incubated in HRP-conjugated secondary antibody against the appropriate host (1:10,000, BioRad) for 1 hour RT. The bands were visualized using Pierce™ ECL Plus western blotting substrate (#32132, Thermo Fisher Scientific). Primary antibodies used were: rabbit anti-phosphorylated TRKB against Y515, Y706 or Y816 (Cell Signaling #4619, #4621, and #4168, respectively); goat anti-TRKB (R&D System, #AF1494) or anti-β-actin mouse monoclonal antibody (Sigma-Aldrich, #A1978). To ensure the quality of the signal, the membranes were stripped only once.

### Protein-fragment Complementation Assay - PCA

Protein-fragment complementation assay (PCA) measures the reconstitution of enzymatic activity of a humanized *Gaussia princeps* luciferase (GLuc) following a direct interaction of the proteins of interest (Kim et al., 2009; Merezhko et al., 2020; Casarotto et al., 2021). Two complementary fragments of the luciferase reporter protein were fused to the intracellular C terminus of TRKB or mutant TRKB.Y433F to produce the wild-type TRKB PCA pair (GLuc1C-TRKB.wt/GLuc2C-TRKB.wt) and the heterozygous TRKB.Y433F PCA pair (GLuc1C-TRKB.wt/GLuc2C-TRKB.Y433F). The GLuc tag was linked via a GS linker that allows the physiological dynamics of TRKB without interference from the presence of the tag. When two TRKB molecules carrying the complementary GLuc fragments dimerize, the reporter refolds in its active conformation thereby producing bioluminescence in the presence of its substrate native coelenterazine. Neuro2A cells, in 10% (v/v) poly-L-Lysine coated white 96 wells (10,000 cells/well) were transfected with the above-mentioned PCA pair constructs. Cells were treated 48h post-transfection with BDNF (10ng/ml/10min), and luminescence measured as a direct indication of TRKB homodimerization with a plate reader (Varioskan Flash, Thermo Scientific, average of 5 measurements, 0.1 s each) immediately after the injection of the coelenterazine substrate (Nanolight Technology).

### Sholl analysis of neurite branching in cortical cultures

Cortical neurons were incubated with BDNF 20 ηg/ml or veh for 15 min once a day for 3 days (7, 8 and 9 DIV in the afternoon). At 9 DIV (in the morning), the neurons were incubated with MgCl_2_ 10 µM for 1h to prevent excitotoxicity and then transfected with mCherry construct, using lipofectamine following manufacturer’ s instructions. The culture medium was replaced by fresh Neurobasal medium supplemented only with L-glu, B27 and penicillin/streptomycin 90 min after the transfection. 24h after the last BDNF treatment, the coverslips were fixed with PFA 4% for 20 min, blocked, and stained with chicken polyclonal anti-MAP2 1:5,000 (#ab5392, Abcam) and Hoescht 1:10,000 for 10 min. Coverslips were washed in PBST (3x) and miliQ water once, then mounted in DAKO fluorescence mounting media (S3023, Dako North America, Inc.). Imaging was acquired in Zeiss LSM700 confocal microscope, 25x oil objective at 1024×1024 pixel resolution, using 647 ηm (for MAP2) and 568 ηm (mCherry) channels. At least 10 Z-stack steps were acquired. Confocal pictures were analyzed in ImageJ (Fiji) software, by compiling the 568 ηm z-stacks, setting the threshold automatically (Shanbhag threshold, available in the software), setting the center of the soma manually, and then counting the branching intersections automatically with built-in Sholl analysis tool (Ferreira et al., 2014). Sholl circle interval was set to 5 µm and the intersections were counted up to 100 µm of distance from the soma.

### Behavioral Analysis

All behavioral trials in the present study were conducted in experimentally naive animals.

#### Cued Fear Conditioning - CuedFC

Male TRKB.Y433F mice and their TRKB.wt littermates were submitted to the fear conditioning protocol to tone followed by two extinction trials, as described previously (Karpova et al., 2011). Briefly, mice were first cue/shock conditioned by co-terminating 5 tone conditioned stimuli (CS of 80dB, 1Hz, 30s) with an unconditioned stimulus foot-shock (US of 0.6mA, 1s) in an inter-trial interval of 20-120s. Over the next 2 days, the freezing-conditioned response was extinguished by subjecting mice to 12 of these same tones (inter-trial interval of 20-60s) each day. Fear conditioning and extinction were performed in different contexts, respectively context A of transparent Plexiglas walls with metal grids on the floor and context B of non-transparent black Plexiglas walls with a flat floor. For each behavioral procedure, mice remained undisturbed in the experimental room for 1h for acclimatization; and mice were allowed to explore the environment for 2min before the presentation of the first tone. The chamber was cleaned with 70% ethanol solution before each animal change. Freezing levels are presented as the sum of all freezing time spent on every tone-trial within each extinction session and were determined by the software (TSE, Bad Homburg, Germany).

#### Context Discrimination Task - CDT

Male TRKB.Y433F mice and their TRKB.wt littermates were submitted to context discrimination task as described previously (Michels et al., 2018; Laukkanen et al., 2021). Briefly, the animals were acclimated to the experimental room for 1h, following 5 min habituation to the experimental chamber, the animals received 3 scrambled shocks (0.6mA/2s, intervals 30s-1min) in context A (conditioning, transparent walls, LxWxH: 23×23×35cm, with metal grid bottom) followed by a 2min period without any shocks. On the following day, the animals were exposed to the unfamiliar context B (black walls with the same dimensions, and black sleek bottom) in a 5 min session. On day 3, the animals returned to the familiar context A for 5 min. The time spent in freezing (s) was determined by the software (TSE, Germany) in the full 5min for each context.

#### Elevated Plus Maze - EPM

Male and female TRKB.Y433F mice and their TRKB.wt littermates were submitted to the elevated plus maze as described previously (Laukkanen et al., 2021). Briefly, the animals were acclimated to the experimental room for 1h, and submitted (5 min session) to an acrylic-built elevated plus-maze. The apparatus was composed of two open arms (30×5cm, a 1cm rim to prevent falls) and two enclosed arms (same dimensions with a 25cm wall). The arms were connected by a central platform (5×5cm) and the apparatus was elevated 40cm above the floor. The percentage of time and entries in the open arms were determined by the software (Ethovision XT 13, Noldus, Netherlands).

### Statistical Analysis

ELISA, Sholl and PCA experiments, as well as CuedFC, and CDT were analyzed by two-way ANOVA followed by Fisher’ s LSD or Tukey’s (for Sholl) when appropriate. WB and EPM data were analyzed by Student’ s t test. All data used in the present study is stored in FigShare under a CC-BY license (DOI:10.6084/m9.figshare.20655042).

## Results

### Direct Binding Assay - DBA

The mutations used in this study are indicated in figure 1A. The Y433 (tyrosine) residue from rat TRKB was mutated to F (phenylalanine), we also substituted the whole TRKB TM domain to the rat TRKA sequence (TRKB/TRKA.TM). As seen in figure 1B, the Y433F mutation or the TRKA transmembrane domain in TRKB do not alter the binding of bBDNF to TRKB. The constant of dissociation (Kd±SEM, pM, n=6/group) observed were: TRKB.wt = 0.3904±0.0877; TRKB.Y433F = 0.4375±0.1069; TRKB/TRKA.TM = 0.3874±0.0747.

**Figure 1.**
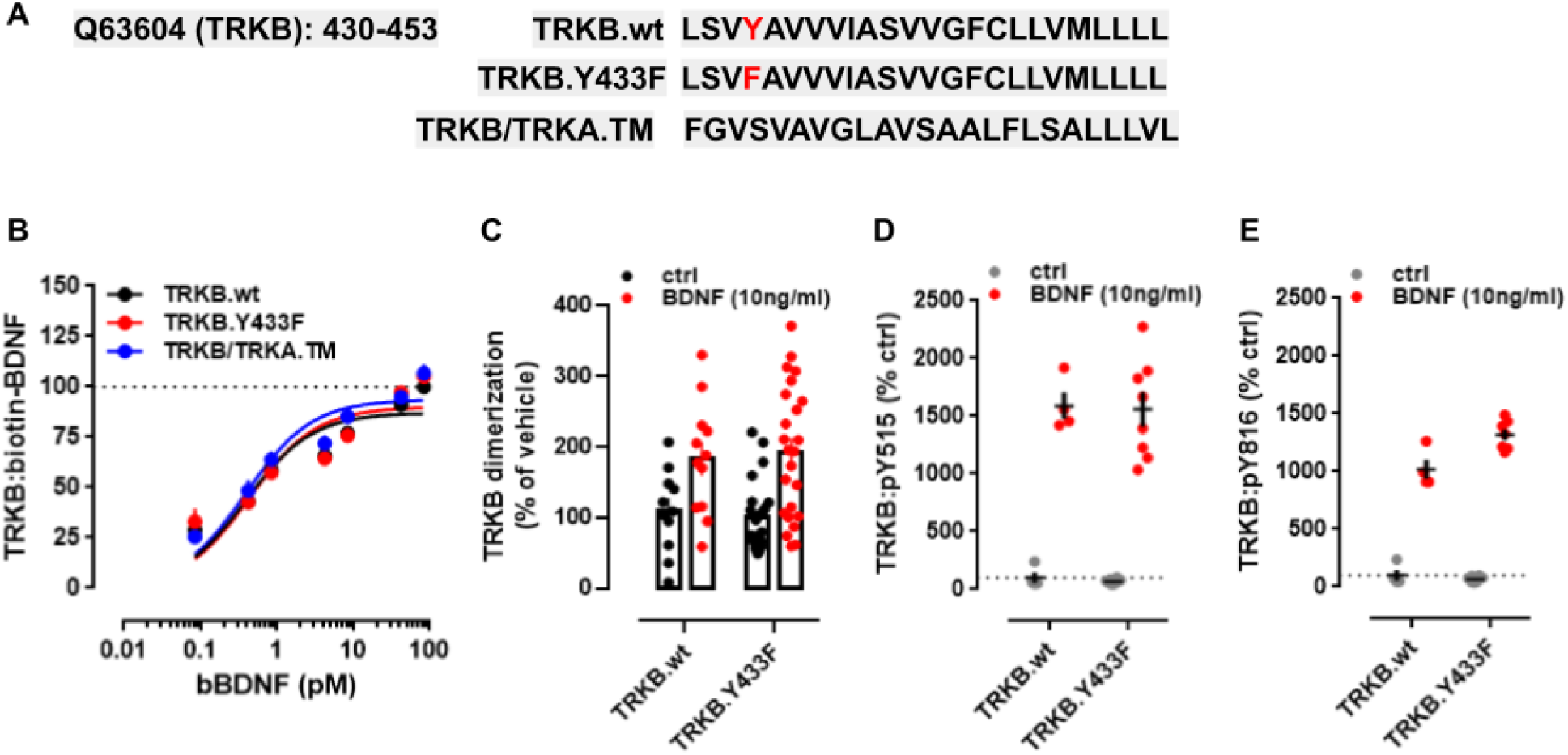
**(A)** The sequence of TRKB transmembrane (TM) domain from rat (UniProt: Q63604, residues 430-453) was mutated at Y433 residue (Y433F) or the entire motif was substituted by the rat sequence of TRKA TM (P35739, residues 419-442). **(B)** The interaction between biotinylated BDNF (bBDNF) and TRKB is not affected by the TRKB.Y433F mutation or in TRKB/TRKA.TM constructs. **(C)** TRKB.Y433F does not prevent the BDNF-induced dimerization of TRKB. The constructs used allow the formation of TRKB.wt homodimer or TRKB.wt/Y433F heterodimer. **(D**,**E)** Analysis of TRKB phosphorylation shows that, cortical cultures from TRKB.Y433F heterozygous mouse embryos respond to BDNF similarly to the the TRKB.wt littermates, regardless of the tyrosine residue tested (**D**: Y515, **E**: Y816).

### Protein-fragment Complementation Assay - PCA

The Y433F mutation in TRKB did not prevent the BDNF-induced dimerization of this receptor as indicated by the two-way ANOVA [interaction: F(1,67)= 0.2658, P=0.6079; genotype: F(1,67)=0.0006266, P=0.9801; BDNF: F(1,67)=20.13, P < 0.0001, N=11-13/group]; figure 1C. As expected BDNF induced an increase in the dimerization of TRKB.wt, and no alteration on BDNF-induced dimerization of the heterodimer TRKB.wt/Y433F was observed.

### *Phospho-TRKB levels in cortical cultures of* TRKB.Y433F *mice*

The effect of BDNF on TRKB phosphorylation (pTRKB) is retained in primary cultures of Y433F mutants (N=4-8/group). No effect of Y433F on BDNF-induced pTRKB levels was observed by two-way ANOVA on pTRKB at Y515 [interaction: F(1,20)=0.002, P=0.9962; genotype: F(1,20)=0.07156, P=0.7918; BDNF: F(1,20)=160.1, P < 0.0001, figure 1D] or at Y816 [interaction: F(1,19)=12.98, P=0.0019; genotype: F(1,19)=8.389, P=0.0093; BDNF: F(1,19)=557.7, P < 0.0001]; figure 1E.

### Neuritogenesis in cortical cultures of TRKB.Y433F mice

BDNF effectively induces neuritogenesis regardless of the genotypes investigated (figure 2). Two-way ANOVA identified effect of the distance from soma (F_18_, 1064= 20.51, p<0.0001) and the groups (TRKB.wt veh, TRKB.Y433F veh, TRKB.wt BDNF, TRKB.Y433F BDNF; F_3,1064_ = 23.79, p<0.0001), but no interaction between those factors (F_54, 1064_= 1.021, p=0.4353). Tukey’ s Multiple Comparison Test pointed to a significant effect of BDNF in the WT mice (veh vs BDNF: p<0.0001) and also in the TRKB.Y433F mutants (veh vs BDNF: p<0.0001). No difference was found when comparing TRKB.wt veh *versus* TRKB.Y433Fveh (p=0.7921), nor TRKB.wt BDNF *versus* TRKB.Y433F BDNF (p=0.3636).

**Figure 2.**
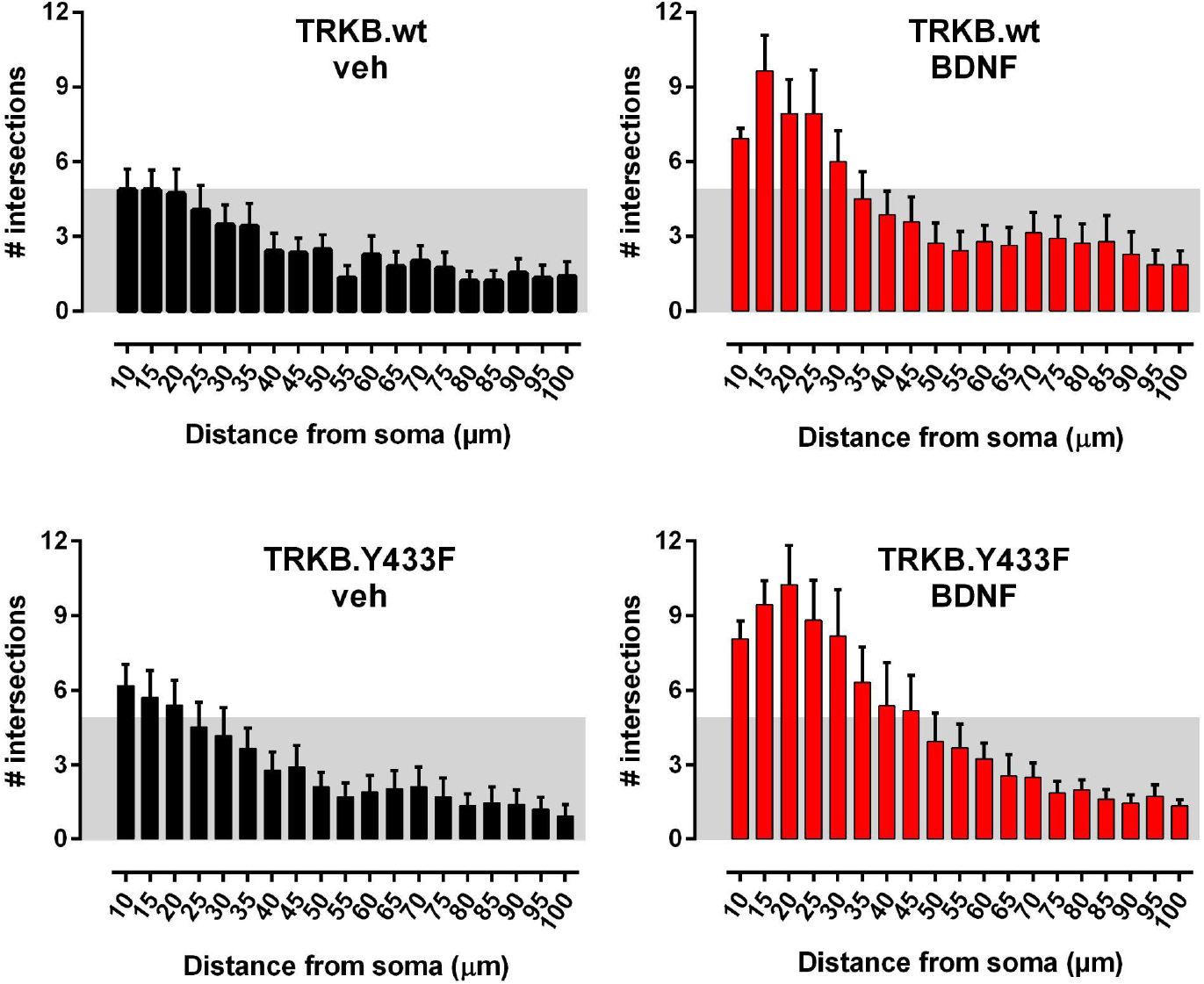
Sholl analysis of the neuritogenesis in cultures from the cortex of TRKB.wt (upper panels) and TRKB.Y433F heterozygous mouse embryos (bottom panels). There was no basal difference between TRKB.wt and TRKB.Y433F heterozygous. Additionally, the neurons respond similarly to BDNF treatment (right panels, in red), regardless of the genotype. Data presented as mean±sem.

### TRKB and pTRKB levels in the prefrontal cortex of TRKB.Y433F adult mice

In adult TRKB.Y433F mice the pTRKB levels measured in the prefrontal cortex are not affected by the genotype [pTRKB Y515: t(12)= 0.2665, P= 0.7944; Y816: t(12)= 0.5650, P= 0.5825; Y706/7: t(12)= 0.6503, P= 0.5277; total TRKB: t(12)= 0.5396, P= 0.5993; beta-actin, ACTB: t(12)=0.3391, P= 0.7404; N=6/group], as seen in figure 3A-E.

**Figure 3.**
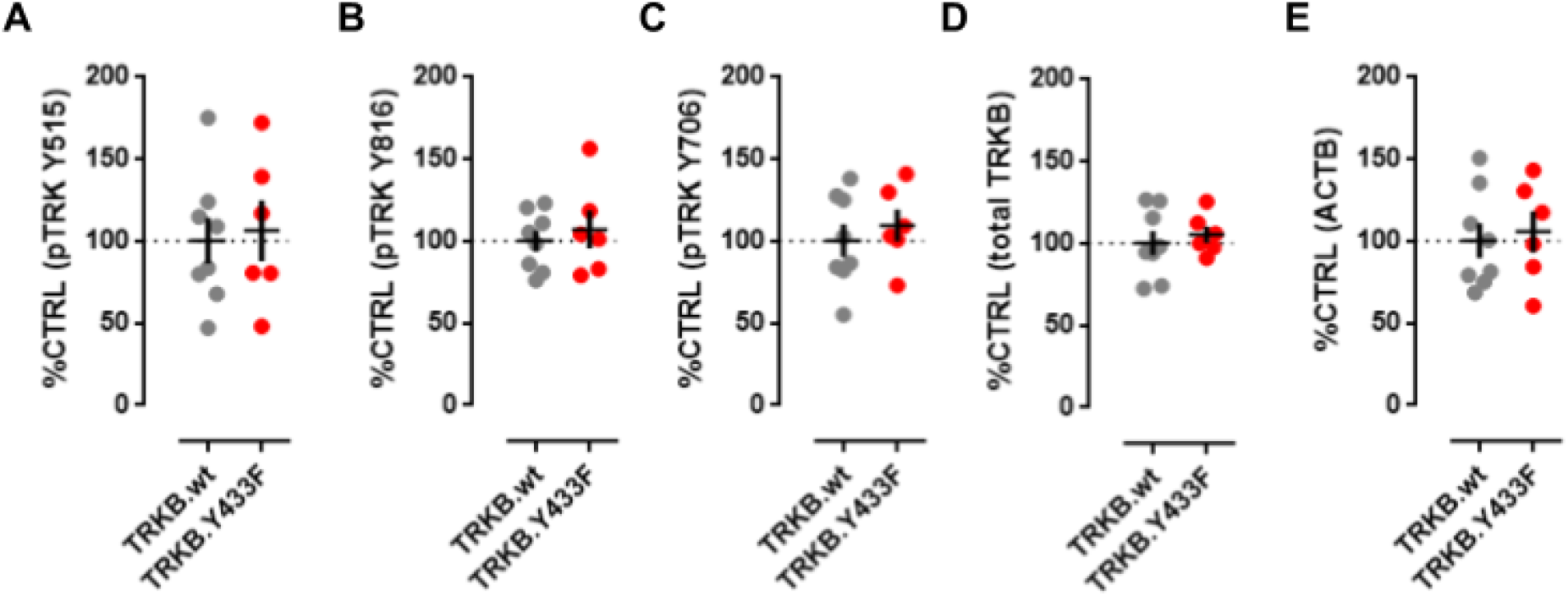
The basal levels of phosphorylated TRKB are not different in the prefrontal cortex of TRKB.wt and TRKB.Y433F mice (A: Y515, B: Y816, C: Y706, D: total TRKB, E: beta actin, ACTB).

### Behavioral analysis of TRKB.Y433F mice

#### Cued Fear Conditioning - CuedFC

No genotype effect of Y433F mutation was observed in the extinction of conditioned fear reaction [interaction: F(1,19)=0.8641, P=0.3643; trial: F(1,19)=7.367, P=0.0138; genotype: F(1,19)=0.2931, P=0.5945, N=10,11], as seen in figure 4A.

**Figure 4.**
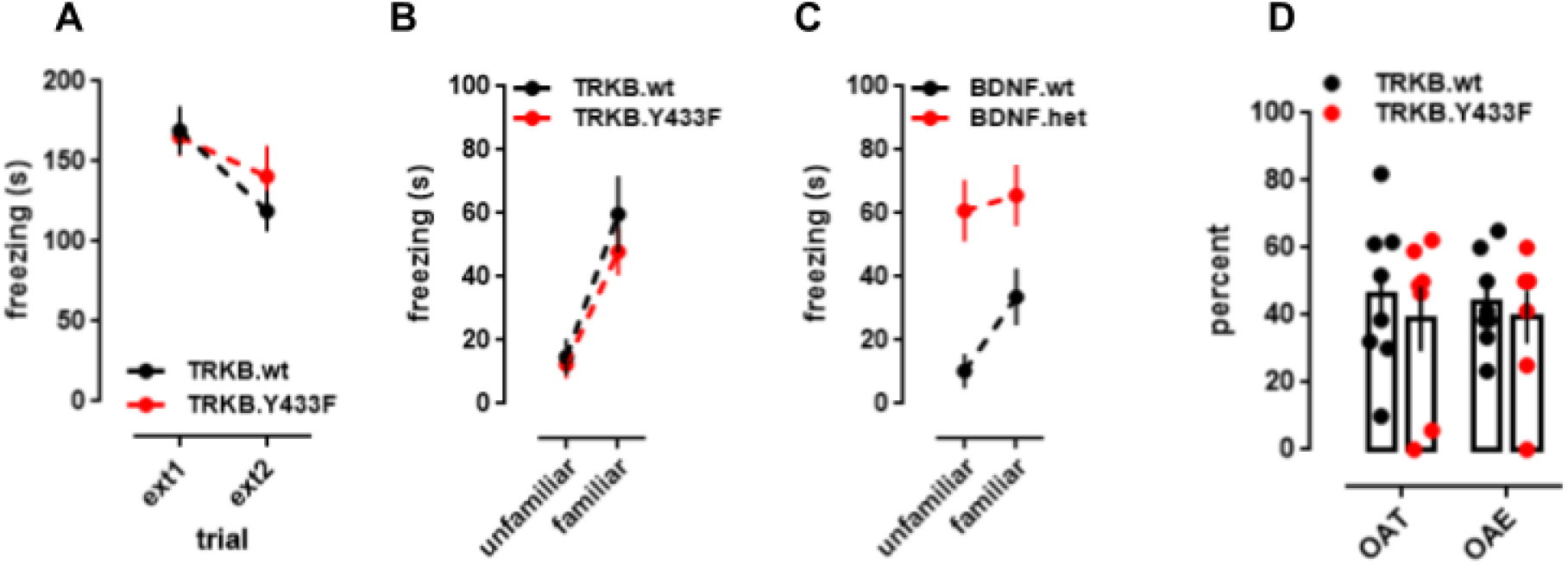
Behavioral Phenotype of TRKB.Y433F mice in fear extinction and elevated plus maze. **(A)** TRKB.Y433F mice exhibit a similar extinction profile of cued fear conditioning response. This strain also did not show any deficits in fear extinction to context [see (Casarotto et al., 2021)]. **(B)** TRKB.Y433F mice are able to discriminate between familiar and unfamiliar contexts previously associated with footshocks. **(C)** For comparison, BDNF.het mice show impaired response to the context discrimination task. **(D)** TRKB.Y433F mice do not show any changes in the anxiety-related parameters of elevated plus maze.

#### Context Discrimination Task - CDT

No genotype effect of Y433F mutation was observed in the ability to discriminate between conditioned and unconditioned contexts [interaction: F(1,15)=0.6472; P=0.4337; trial: F(1,15)=45.87; P < 0.0001; genotype: F(1,15)=0.5711, P=0.4615; N=8,9], figure 4B. For comparison, BDNF.het mice have compromised performance in this test [interaction F(1,10)=6.745, P=0.0266; trial: F(1,10)=15.64, P=0.0027; genotype: F(1,10)=12.57, P=0.0053; N=6/group], figure 4C, as observed previously (Laukkanen et al., 2021), as well as in the extinction of conditioned fear (Karpova et al., 2011).

#### Elevated Plus Maze - EPM

as seen in figure 4D, no effect of the genotype was observed in the performance on the EPM [percent time in the open arm, %OAT: t(13)=0.5657, P=0.5813; percent entries in the open arm, %OAE: t(13)=0.4779, P=0.6407, N=8,7].

## Discussion

Our findings demonstrate that hTRKB.Y433F mutation has no effects on the baseline or BDNF-induced TRKB signaling, or on TRKB-mediated behavior. This confirms our previous findings that the hTRKB.Y433F mutation has no effect on basal TRKB signaling, TRKB plasma membrane localization, and on basal behavior of hTRKB.Y433F mutant mice (Casarotto et al., 2021). In sharp contrast, essentially all the effects of antidepressants and ketamine were lost in these mice (Casarotto et al., 2021). These data suggest that this heterozygous mutation selectively inhibits the effects of drugs that directly bind to the TRKB CARC domain without having any effects on basic or BDNF-induced TRKB signaling, which makes this mouse an excellent experimental model for the antidepressant research.

Binding of BDNF to TRKB.Y433F was indistinguishable from that of the wildtype TRKB. Moreover, substitution of the entire TMD of TRKB by that of TRKA also had no effects of BDNF binding. TRKA does not have the ^439^AXXXG^443^ dimerization motif and its TMD is also in other ways divergent from that of TRKB (Nikoletopoulou et al., 2010), which suggests that the TMD configuration does not influence BDNF binding. BDNF binds to the second Ig-domain of TRKB (Schneider and Schweiger, 1991) and this binding is apparently independent of the nature of the TMD.

We have used split luciferase-based protein complementation assay to investigate the effects of the TRKB.Y433F mutation on TRKB dimerization (Remy and Michnick, 2006). Cells co-expressed the TRKB.Y433F mutant and the wildtype TRKB, each tagged with a complementary half the luciferase reporter. Therefore, dimerization that is revealed in this assay can only occur in a heterozygous configuration, between a mutant and a wildtype TRKB. We found that BDNF-induced dimerization is intact with this assay. We previously found that TRKB dimerization was reduced, although not completely prevented, in a homozygous configuration where two TRKB.Y433F mutants carrying complementary fragments of the luciferase were overexpressed (Casarotto et al., 2021).

Cultured cortical neurons derived from hTRKB.Y433F mice showed no differences in the baseline or BDNF-induced TRKB autophosphorylation in response to BDNF administration. Our previous studies showed that when the TRKB.Y433F construct is overexpressed in cultured cells, the response to BDNF stimulation is compromised (Casarotto et al., 2021). Specifically, autophosphorylation of the Y816, a recognition site for PLCγ, is reduced when TRKB.Y433F is overexpressed, while autophosphorylation of the Y515, the shc recognition site, is not compromised. Here we have found that both of these responses are normal in neurons derived from the hTRKB.Y433F mice and the baseline phosphorylation of both these sites is also normal in brain tissue from hTRKB.Y433F mice. Taken together, these findings suggest that for the baseline activity and BDNF responses, TRKB.Y433F mutation acts as a recessive mutation, but for responses to antidepressant drugs, it acts as a dominant, apparent already when a single allele of TRKB is mutated. This conclusion was also supported by the behavioral findings with the hTRKB.Y433F mice. While these mice exhibit a dramatically reduced responsiveness to antidepressants (Casarotto et al., 2021), they showed normal behavior in tests that are known to be sensitive to BDNF signaling. In contrast, mice heterozygous for BDNF null allele show clear behavioural deficits in the context discrimination test, confirming that this response requires intact BDNF signaling.

An exome sequencing study of 197 unrelated patients with developmental and epileptic encephalopathy identified four patients with a similar phenotype and a Y433C (Y434C in human nomenclature) *de novo* point mutation in the TRKB gene, in the very same tyrosine residue that has been mutated to phenylalanine in our studies (Hamdan et al., 2017). While more research is needed to elucidate the exact mechanism of action of this Y433C mutation, it is possible that the developmental and epileptic encephalopathy is produced by a constitutive active TRKB through a disulfide bridge between the mutant cysteine residues. Being a *de novo* mutation and therefore heterozygous, a pair of mutant alleles that can create a S-S bridge is only out of four possible TRKB dimers, nevertheless, only a minority of TRKB dimers when constitutively active may produce a dramatic phenotype. Since mutations of a tyrosine to both cysteine and phenylalanine require a point mutation in only a single nucleotide (A>G and A>U, respectively) and several Y433C mutations have been detected, it is likely that also the Y433F mutation (or Y434F in human TRB) has occurred, however, we have not found that mutation reported in repositories. Our current data indicate that a human TRKB.Y434F mutation would likely be recessive and not produce any overt phenotype as heterozygous, which may explain why it has not been detected so far. Nevertheless, these findings highlight the importance of the tyrosine-433 (434 in case of humans) for TRKB signaling.

Taken together, we have found that a hTRKB.Y433F mutation does not interfere with the baseline TRKB function or with BDNF signaling and it does not produce any phenotype in behavioral tests that are known to be sensitive to BDNF signaling. However, plasticity-promoting and antidepressant-like responses to antidepressant drugs that bind to TRKB at the site including the tyrosine-433 are severely compromised in the hTRKB.Y433F mice, indicating that while this mutation is recessive for BDNF responses, it is dominant for antidepressant binding. The hTRKB.Y433F mouse could therefore be of substantial interest in the process of screening for novel potential antidepressants or other drugs that promote plasticity by binding to the TRKB transmembrane domain.

## Acknowledgements

The authors thank Seija Lågas and Sulo Kolehmainen, and the LAC (laboratory animal center) personnel Vootele Voikar and Neli Koivisto, for their technical assistance. This research was funded by the ERC grant – iPLASTICITY (#322742), the Sigrid Jusélius Foundation, Jane & Aatos Erkko Foundation, and Academy of Finland grants (#307416, #327192, # 347358).

